# The nucleus serves as the pacemaker for the cell cycle

**DOI:** 10.1101/2020.06.16.153437

**Authors:** O. Afanzar, G. K Buss, T. Stearns, J. E. Ferrell

**Affiliations:** Dept. of Chemical and Systems Biology, Stanford Medicine, Stanford CA 94305-5174; Dept. of Molecular and Cellular Physiology, Stanford Medicine, Stanford CA 94305; Dept. of Biology, Stanford University, Stanford CA 94305; Dept. of Genetics, Stanford Medicine, Stanford CA 94305; Dept. of Biochemistry, Stanford Medicine, Stanford CA 94305

## Abstract

Mitosis is a dramatic cellular process that affects all parts of the cell. In *Xenopus* embryos and extracts it is driven by the activation of a bistable trigger circuit, whose various components are localized in the nucleus, centrosome, and cytoplasm. In principle, whichever cellular location has the fastest intrinsic rhythm should act as a pacemaker for the process. Here we followed tubulin polymerization and depolymerization in *Xenopus* egg extracts supplemented with demembranated sperm, and thereby identified locations where mitosis first occurred. We found that mitosis was commonly first initiated at sperm-derived nuclei and their accompanying centrosomes, and then spread outward in circular trigger waves. The cell cycle was ∼20% more rapid at the nucleus/centrosome-associated trigger wave sources than in the regions of the extract that appeared not to be entrained by trigger waves. Nuclei produced from phage DNA, which did not possess centrosomes, also acted as trigger wave sources, but purified centrosomes in the absence of nuclei did not. We conclude that the nucleus accelerates mitotic entry and propose that it acts as a pacemaker for cell cycle.

**One Sentence Summary:** Studies in cycling *Xenopus* egg extracts show that mitosis first occurs in the nucleus and then spreads outward through the cytoplasm in circular trigger waves.

---

Mitotic entry is driven by a circuit of proteins that regulates cyclin-dependent protein kinase-1 (Cdk1) and its opposing phosphatases. The circuit includes at least 5 interlinked positive and double-negative feedback loops and functions as a bistable toggle switch, which transitions from a stable interphase state with low Cdk1 activity and high PP2A-B55 activity, to a stable M-phase state with high Cdk1 activity and low PP2A-B55 activity (*1-5*). In some contexts, like the *Xenopus laevis* embryonic cell cycle, the cell cycle operates as a relaxation oscillator (*6*), with the bistable triggering firing, and the cell cycle repeating, with a precise (± a few percent) (*7-9*) fixed period.

Systems with a bistable trigger have the potential to generate trigger waves, propagating fronts of activity that spread outward from a source without slowing down or diminishing in amplitude (*10, 11*). Familiar examples of biological trigger waves include the action potential (*12*), calcium waves (*13-15*), and apoptotic waves (*16*), and, as predicted in the 1990s (*17*), mitosis does spread through the cytoplasm via constant velocity (∼60 µm/min) trigger waves (*18*).

A trigger wave typically originates from wherever the oscillator has the fastest intrinsic rhythm. For example, heartbeats originate at the sinoatrial node, which has a typical frequency of 60-90 min^−1^ in humans, and spread to and override the slower rhythms of the atrioventricular node (whose intrinsic frequency is 40-60 min^−1^) and ventricles (30-40 min^−1^) (*19*).

At present it is uncertain where mitotic trigger waves begin. Plausible candidates include the centrosome, where the cyclin B1-Cdk1 and Cdc25C have been found to concentrate in late interphase (*20, 21*); the nucleus, to which cyclin B1-Cdk1 and Cdc25C translocate just prior to prometaphase (*22-24*); or the cytoplasm, since cyclin A2-Cdk1/2 may shift from the nucleus to the cytoplasm late in interphase (*25-28*). However, studies with FRET probes have suggested that perhaps mitosis is initiated in all parts of the cell essentially simultaneously (*29*).

Here we set out to determine what subcellular compartment serves as the pacemaker for mitotic oscillations.

## Results

We made use of *Xenopus* egg extracts, living cytoplasm that can carry out the biochemical reactions of the cell cycle (*30*), into which we added various candidate pacemaker structures. We placed small volumes (3 µl) of extract supplemented SiR-tubulin (a fluorogenic dye that binds polymerized tubulin), and GST-NLS-mCherry (a nuclear marker to monitor the breakdown of the nucleus at the onset of mitosis) in the wells of a 96-well plate under mineral oil (Fig. 1A), and monitored cell cycle dynamics in this essentially two-dimensional system by time-lapse video microscopy.

**Fig. 1.**
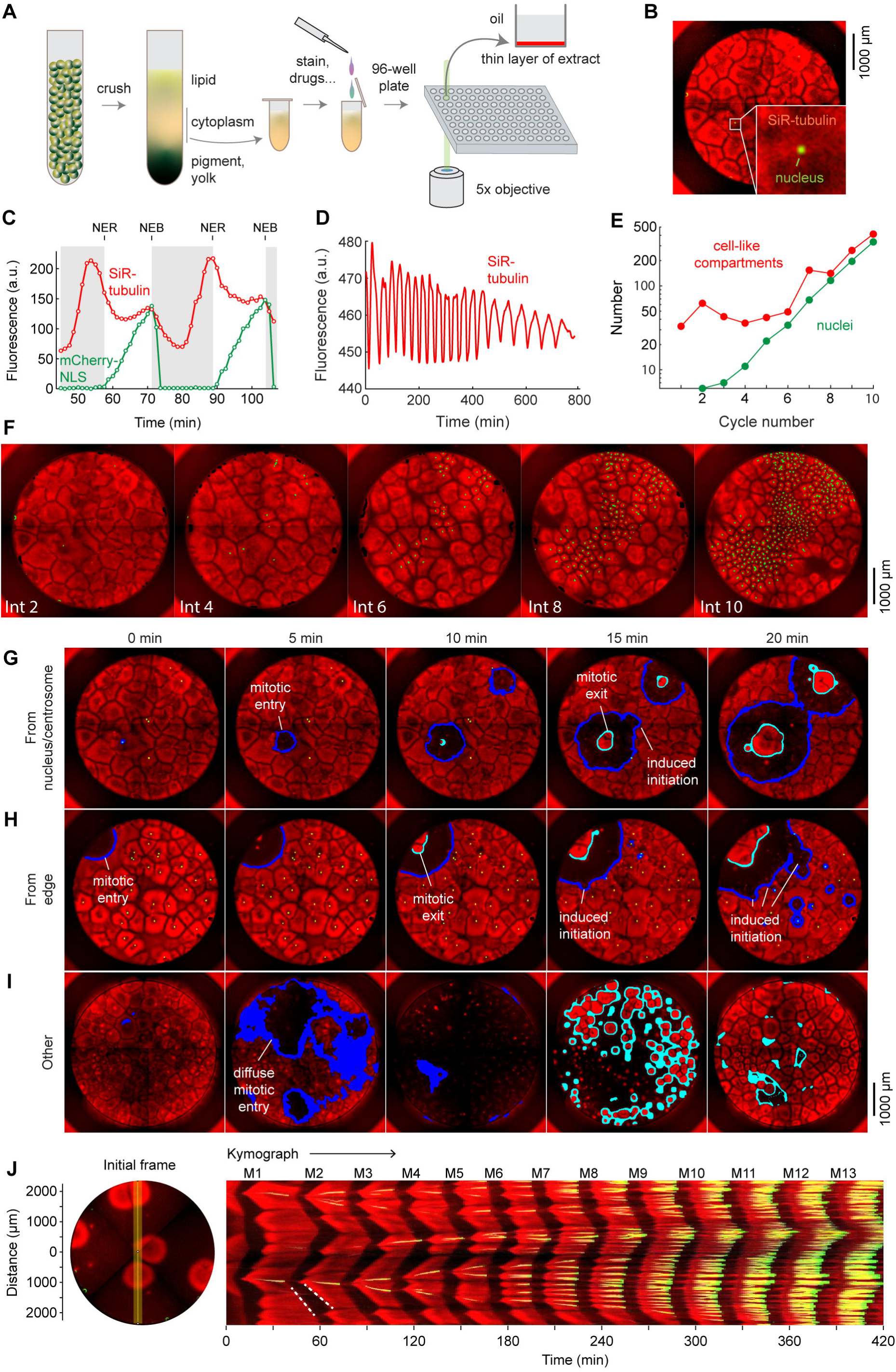
Self-organization, robust cycling, and two-dimensional trigger waves in extracts supplemented with demembranated sperm chromatin. (A) Experimental scheme. (B) A snapshot of interphase 2 from one experiment. A nucleus is shown in green (also shown in enlargement) and microtubules are in red. (C) Normalized fluorescent intensity for the nucleus and microtubules from the area shown in the enlarged area of B, as a function of time. (D) Periodic microtubule fluorescence over the entire time course of the experiment. (E) The number of nuclei and cell-like compartments as a function of cycle number from the same experiment as in B. (F) Cell-like compartments and nuclei through ten cycle cycles. (G-I) Types of mitotic sources. (G) Trigger waves emanating from nuclei and/or their accompanying centrosomes. (H) Trigger waves originating at the edge of the well. (J) Other sources. In the example shown here, mitosis begins in many regions of the plate nearly simultaneously. (J) A kymograph (right) from a cross section an extract over 14 mitotic cycles. The dashed white line show linear spread of microtubule depolymerization at the onset of mitosis, with a speed of 65 µm/min, and aster growth during the transition back to interphase, with a speed of 40 µm/min.

### Continuous monitoring of mitotic entry and exit

Initially we included small concentrations (< 10 sperm/mm^2^) of demembranated sperm chromatin in the extracts. The sperm chromatin recruits bits of endoplasmic reticulum to form functional nuclei, and the also provide associated centrosomes, which promote microtubule organization. As previously reported (*31*), upon warming, the extracts began to self-organize into cell-like compartments. Typically several such compartments were present in the first interphase, with sperm nuclei near the centers of the best organized compartments (e.g. movie S1, Fig. 1B and J). Generally by the second interphase the whole well of cytoplasm was organized into compartments.

The extracts underwent regular, periodic cycles of nuclear envelope breakdown and reformation, accompanied by cycles of microtubule depolymerization and repolymerization (movie S1, Fig. 1C). The extracts typically continued to cycle for ∼12 hours (movie S1, Fig. 1D). There was an exponential increase in the number of nuclei over this time period, although the fold-increase per cycle was typically less than a full doubling (Fig. 1E-F) (1.71 ± 0.01 fold for the experiment shown here). The number of cell-like compartments also grew exponentially, but only after a time lag (Fig. 1E). This is because typically only the compartments that contained nuclei and centrosomes divided, and initially that was a small proportion of the compartments.

Mitosis often began at or near a nucleus and its accompanying centrosomes (Fig. 1G, movie S2). Waves of mitotic initiation and microtubule depolymerization then propagated outward from these sources as circular fronts that expanded at constant speed. As the nuclei and centrosomes0020replicated and divided, they often produced new trigger wave sources centered on the daughter nuclei and centrosomes (movie S3).

Mitotic waves also originated at discrete locations close to the edge of the well (Fig. 1H, movie S2). This edge effect has been noted previously in studies of mitotic trigger waves and apoptotic trigger waves in *Xenopus* extracts (*16, 32*), and it is seen in mathematical models of trigger waves as well (*33*). Mitosis also sometimes initiated from locations that were neither close to nuclei nor to the edge of the well (denoted “others” in Fig. 1I, movie S2). These sources were often diffuse, with mitosis apparently beginning at many locations nearly simultaneously and quickly engulfing large regions of cytoplasm (Fig. 1I). These “other” wave sources were often seen in an extract’s first cell cycle, and sometimes near the end of mitosis in later cycles.

Because the speeds of the irregular, ill-defined waves that appeared to spread from these sources were often much higher than those of bona fide mitotic trigger waves, they most likely represent phase waves—a type of wave that arises from differences in intrinsic timing rather than spatial coupling, like the waves of flashing lights on a movie marquee (*10, 34*).

The mitotic wave fronts are highlighted in blue in Fig. 1G-I, and the areas where microtubules are reappearing as the cytoplasm transitions to the next interphase are shown in cyan. Often the mitotic waves advanced smoothly from one cell-like unit to the next, but sometimes the front appeared to induce a nearby source to fire a trigger wave, as seen at the 15 min time points in Fig. 1G and H.

### Quantitative characterization of the two-dimensional trigger waves and their sources

We analyzed the speeds of the mitotic trigger waves by plotting kymographs; a constant-velocity mitotic wave emanating from a well-defined source can be seen as a straight line of disappearing microtubule fluorescence (Fig. 1J). For example, at the start of mitosis 2 in this experiment, waves of microtubule (red) depolymerization originated in the vicinity of the two nuclei (green) and spread outward at a speed of 65 µm/min, similar to previously-reported mitotic wave speeds (*18*). This was followed about 10 min later by a slower (40 µm/min) wave of microtubule polymerization that spread out from a microtubule aster. This velocity is about a factor of two higher than that reported by Ishihara and colleagues (*35*).

To characterize and analyze mitotic dynamics more comprehensively and in the full two-dimensional context, we used an automated approach to identify, map, and characterize mitotic sources from time-lapse video data from 24 experiments. Mitotic waves were taken to be fronts of microtubule depolymerization that expanded from one frame to the next. For the example shown in Fig. 2A, at the 51 min time point, 11 such waves were identified. Note that in this cycle, initially (at 46 min) there were numerous microtubule-depleted areas that superficially resembled mitotic regions but did not expand with time, and so were not classified as mitotic. The centroid of a mitotic region in its first frame was generally taken to be the mitotic source. There were a few instances where the region was dumbbell-shaped; in these cases we considered mitosis to have originated nearly simultaneously from two distinct sources.

**Fig. 2.**
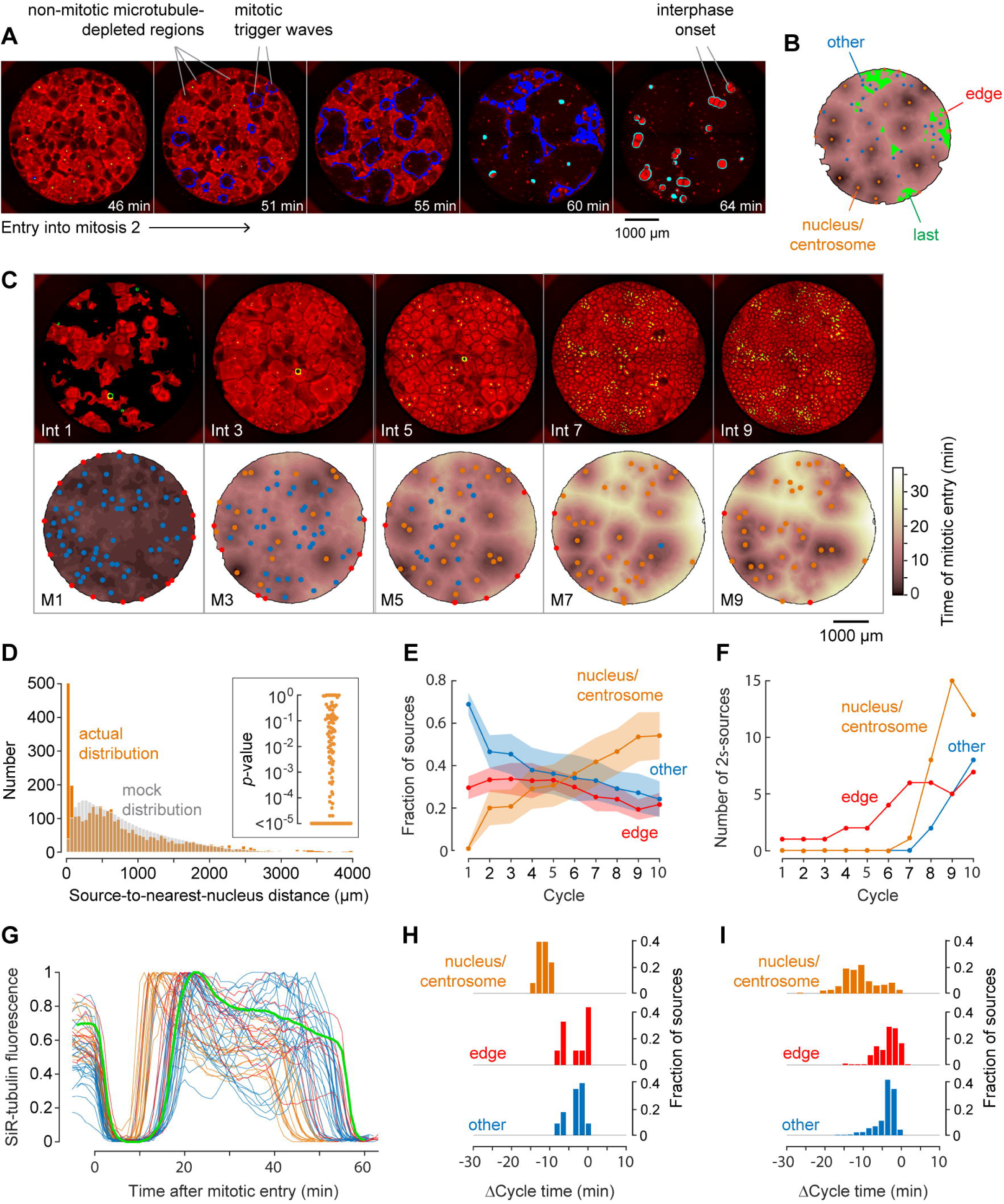
Demembranated sperm promotes mitotic initiation by shortening cycle time. (A) Typical sperm-mediated mitotic trigger waves. Nuclei (NLS-mCherry) are shown in green and microtubules (SiR-tubulin) in red. The advancing fronts of mitosis are depicted in blue and the advancing fronts of interphase in cyan. (B) The trigger wave sources for the same cell cycle shown in (A), identified and characterized automatically. The time at which region of the extract entered mitosis is depicted by a color scale, ranging from 46 min (dark) to 64 min (light). The latest 15% of the cytoplasm to enter mitosis is color-coded green. The trigger wave sources are plotted as filled circles of various color: orange for sources associated with nuclei/centrosomes, red for sources at the edge of the well, and blue for others. (C) Montage showing changes in the pattern of mitotic sources from cycle 1 to 9. The coloring denotes the time at which each pixel entered mitosis relative to the earliest pixels in that cycle. (D) The distance of the trigger wave sources to the nearest nucleus (orange), compared to a mock distribution (gray). Data are from 109 cell cycles from 24 experiments and include 3402 sources. The inset shows the probabilities (*p*-values) of obtaining the observed number of mitotic sources (or more than the observed number) that are close (< 100 µm) to nuclei for the 109 cell cycles analyzed individually. *p*-values were calculated by bootstrapping with 10^5^ randomized source positions for each individual cycle. (E) The fraction of the mitotic sources classified as associated with nuclei/centrosomes (orange), edges (red), or others (blue) as a function of cycle number. The data points are mean fractions from 10 experiments where the extracts cycled at least 10 times, and the shaded regions show the ± SEM. (F) The number of sources whose time of initiation was more than two standard deviations earlier than the average sources of the same experiment. (G) Time courses of microtubule fluorescence intensity, normalized to the maximum and minimum fluorescence in that cycle and time-aligned to make mitotic initiation occur at t = 0. The initiation of mitosis was taken as the time when the microtubule fluorescence first showed a significant decline. (H) Acceleration of the cell cycle, as measured by Δcycle time, at the different cell cycle sources relative to the slowest 15% of the pixels for the experiment shown in (G). The Δcycle times were calculated as the time between the onset of the mitosis after nuclei appeared and the onset of the preceding mitosis, relative to the slowest 15% of the well. (I) Histograms of Δcycle times relative to the slowest 15% of the pixels for mitotic sources from 21 experiments. The total numbers of sources are 188 nucleus/centrosome sources, 195 edge sources, and 420 others.

We then constructed a map of the mitotic sources and the mitotic dynamics of the cell cycle under consideration. We colored the pixels between the sources based on the times at which they entered mitosis. For the cell cycle shown as a montage in Fig. 2A, the resulting map is shown in Fig. 2B. It can be regarded as a topographic map, with sources sitting at the bottoms of dark valleys of various depths; the deeper the valley, the earlier the time at which mitosis happened at that source. In the experiment shown in Fig. 2A and B, the sources associated with nuclei/centrosomes (orange) were deep and controlled relatively large areas of cytoplasm, whereas the “other” (blue) sources and the single edge-associated source (red) were shallow and controlled little cytoplasm. The last 15% of the pixels to enter mitosis, which often were regions that appeared not to have been entrained by a trigger wave, were also identified (Fig. 2B, green). We used these late pixels as a benchmark, and expressed the time at which mitotic entry was advanced at the various trigger wave sources relative to these late regions.

The evolution of the sources and mitotic dynamics through the first 9 cycles of the experiment used for Fig. 2A and B is shown in montage form in Fig. 2C and in movie S4. In the first cycle, most of the extract entered mitosis at about the same time. Numerous mitotic sources were identified, mostly in the “other” (blue) category, and the sources were shallow. By the third cycle, nuclei had appeared, and many of them acted as sources for trigger waves. By the ninth cycle, the depths of the deepest nuclear trigger wave sources had increased, most of “other” sources had disappeared.

We used the automated procedure to map the mitotic sources in 109 cycles from 24 different experiments. Overall, 700/3402 (20%) of the sources were close (< 100 µm) to nuclei and their accompanying centrosomes. This proportion was ∼5-times higher than would be expected by random chance (Fig. 2D). To further test the robustness of this association, we calculated *p*-values by bootstrapping for each of the cycles individually. In 69/109 of the cycles, the number of trigger wave sources close (< 100 µm) to a nucleus was greater than would be expected by chance (*p* < 0.05) (inset, Fig. 2D), supporting the significance of the association between trigger wave sources and nuclei/centrosomes. As the cell cycles proceeded, the proportion of the trigger waves that emanated from nuclei/centrosomes rose (as did the number of nuclei), and the proportion not associated with any obvious structure fell (Fig. 2E). The “best” trigger wave sources—those more than two standard deviations earlier than the average sources, which were also the sources that controlled the largest areas of cytoplasm—tended to be found in the latest cycles (Fig. 2F).

By definition, mitotic sources are regions where the cell cycle is faster than it is in the surrounding cytoplasm. One way to estimate how much faster the cycles are is to look at early cycles where trigger waves are present in some of the well, but have not taken over the whole well, and compare the timing of the cell cycle at the trigger wave sources to that at the slower, unentrained regions. For example, in the kymograph shown in Fig. 1J, at the onset of mitosis 2 there are two trigger wave sources, one near the top of the kymograph and one approximately 800 µm below the middle of the kymograph. The time from the start of M1 to the start of M2 at the two trigger wave sources is 35 and 39 min, respectively, whereas the slowest regions in the middle and at the bottom of the kymograph have cycles of 46 and 48 min, indicating that the cell cycle is on the order of ∼10 min or ∼20% faster at the nuclear trigger wave source than in these slowest region.

The same type of analysis can be extended from the one-dimensional kymograph slice to the whole two-dimensional area of a well using the topographic map representation. For the cycle shown in Fig. 2A and B (a different experiment from the one shown in Fig. 1J), the time courses of the changes in tubulin intensity at each of the mitotic sources and in the benchmark unentrained region are shown in Fig. 2G. The latest regions of the cytoplasm for this experiment had an average cell cycle duration of 54.0 ± 0.6 min (mean ± S.D). Mitotic sources that were in proximity to nuclei entered mitosis 13.4 ± 1.8 min earlier, whereas most of the “other” sources and the edge-associated sources were advanced by lesser amounts (4.5 ± 2.7 min and 4.7 ± 3.9 min, respectively) (Fig. 2G, H). Thus, the nuclei and/or centrosomes appeared to be the most effective pacemakers, and they accelerated the cell cycle by 25 ± 3% in this experiment.

The same trend was seen in the aggregated data from 21 experiments (Fig. 2I), with mitosis beginning from the nucleus-associated trigger wave sources 11.7 ± 4.7 min early (or 19.4% ± 10.0%, mean ± S.D.), and with the other classes of sources being accelerated by smaller amounts (3.8 ± 3.5 min for edge-associated sources and 3.8 ± 3.1 min for other sources, mean ± S.D.).

These findings indicate that either the nucleus, the associated centrosomes, or the two structures together are particularly effective at accelerating the cell cycle and acting as a pacemaker structure. All told, 80% of the sources that accelerated mitosis by more than 10 min were nucleus-associated.

### With no added sperm, extracts cycle and generate trigger waves, but the trigger wave sources have only modestly accelerated cell cycles

Next we examined extracts with no added sperm chromatin. The cell-like compartments formed more slowly than they did in extracts supplemented with sperm (cf. Fig. 1J and 3A), as previously noted (*31*), and the compartments generally did not divide, although in some experiments regions of robust compartment division emerged from one or more locations at the edge of the well during the later cycles (Supplemental Figure S1).

A typical experiment is shown in montage form in Fig. 3A and in movie S5. The first mitosis occurred at all locations nearly simultaneously, although a few relatively shallow edge-associated sources and other sources were identified (Fig. 3A). By the second cycle, two relatively strong edge sources had emerged (at 12:00 and 1:00 on the heat map), and they grew to dominate the mitotic dynamics over the next several cycles. The self-organizing character of trigger waves (*11*) is particularly apparent in these experiments where new sources are not being generated from nuclear division. A kymograph bisecting the stronger of the two sources showed a wave of microtubule depolymerization spreading at a constant speed of 60 µm/min (Fig. 3B), similar to the speeds seen in the presence of sperm.

**Fig. 3.**
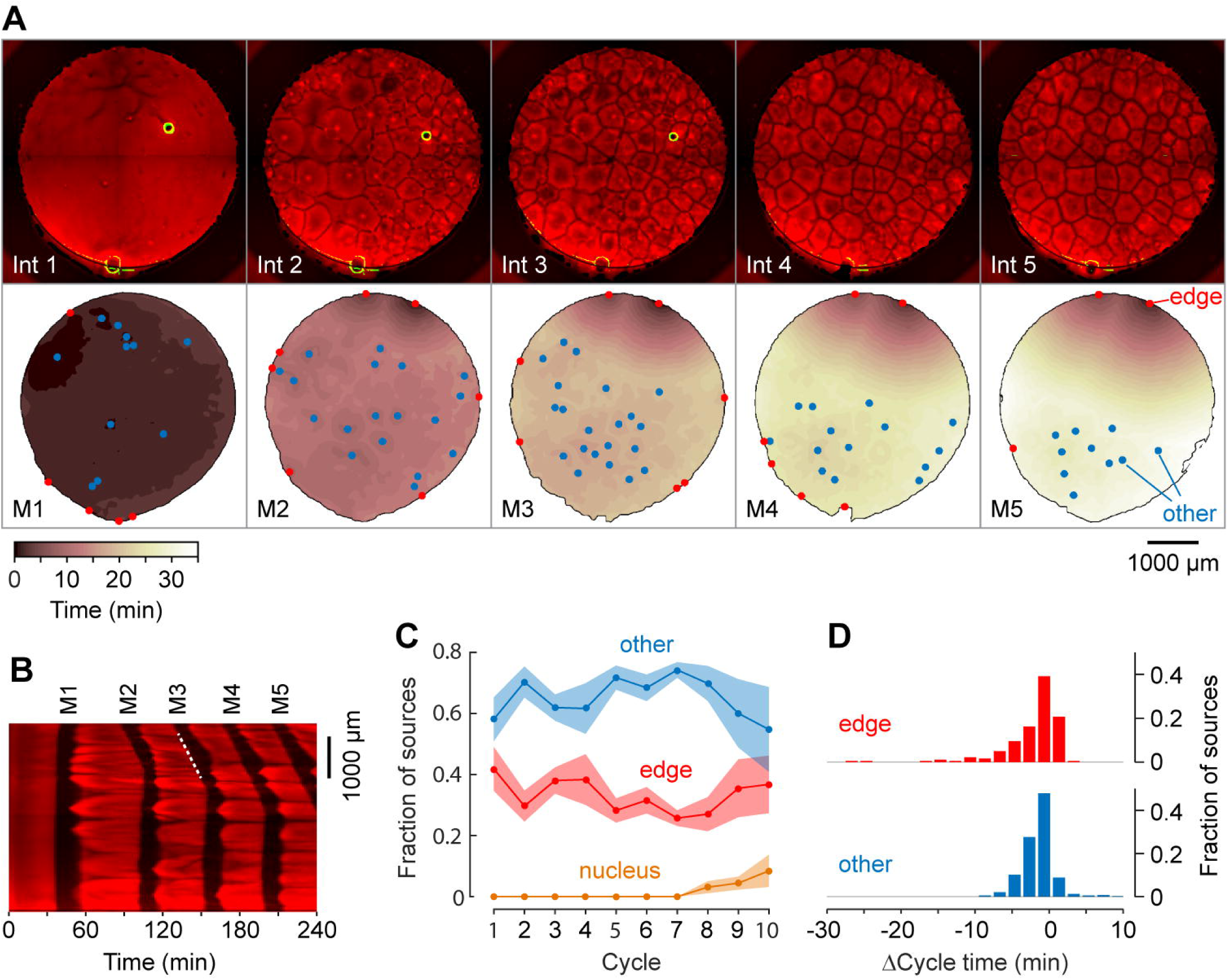
Mitotic waves in cycling extracts with no added sperm chromatin, DNA, or centrosomes. (A) Montage from a typical experiment showing the microtubules (top) and the heat map representation of the mitotic sources (bottom) for the first five cycles. Edge-associated sources are shown in red; other sources—i.e. those not associated with edges or nuclei—are shown in blue (other). The greenish circle in the upper right-hand quadrant of the first three frames of the microtubule montage is a bubble, not a nucleus. The coloring of the heat map representation denotes the time at which each pixel entered mitosis relative to the earliest pixels in that cycle. See also movie S5. (B) Kymograph based on movie S5. The kymograph was calculated for a slice beginning at the edge source at 1:00 in (A) and transecting the well. The trigger wave speed at the entrance to M3 was 60 µm/min (dashed white line). (C) Mean fraction of the mitotic sources originating from edges, nuclei, and other locations as a function of cycle number. Data are from 6 experiments. (D) Acceleration of the cell cycle at edge sources and other sources relative to the slowest 15% of the well. Data are from 23 experiments and include 170 edge sources and 385 other sources.

From six independent experiments, most of the sources were either associated with the edge of the well or with no obvious structure (Fig. 3C). In some experiments, nuclei began to appear and replicate in the later cycles even though no sperm had been added (Fig. 3D), presumably as a result of replication of small amounts of egg DNA present in the extracts. Most of the sources appeared to accelerate the cell cycle by very little; for the edge sources the average Δcycle time was −3.1 ± 5.3 min, with 10/555 (1.8%) of the edge sources, including the two dominating sources in Fig. 3A, having Δcycle times of 10 min or more (Fig. 3C). Overall the “other” sources had an average Δcycle time of −2.3 ± 2.5 min, with only 1/555 (0.2%) having Δcycle times of 10 min or more (Fig. 3C). Thus, extracts containing no nuclei or centrosomes cycled and generated mitotic trigger waves, but their mitotic sources were generally poorer than nucleus/centrosome-associated sources.

### Centrosomes promote cytoplasmic organization and division, but do not appear to act as trigger wave sources or accelerate the cell cycle

Sperm provide the extract with two plausible mitotic sources, centrosomes and nuclei. To determine whether centrosomes are sufficient to pace the cytoplasm, we added purified HeLa cell centrosomes to the extracts at concentrations of up to 8 centrosomes/mm^2^ and assessed their effect on mitotic dynamics. The added centrosomes produced foci of microtubules during interphase that promoted the rapid organization of cell-like compartments (Fig. 4A, B). The extracts then cycled (shown in montage for in Fig. 4C and in movie S6), with the centrosomes increasing in number during each cell cycle (Fig. 4C) and the cell-like compartments dividing as they did in the presence of sperm (Fig. 4B). Boiled centrosomes did not promote the organization of cell-like compartments, did not replicate, and did not promote the division of the cell-like compartments (Supplementary Figure S2).

**Fig. 4.**
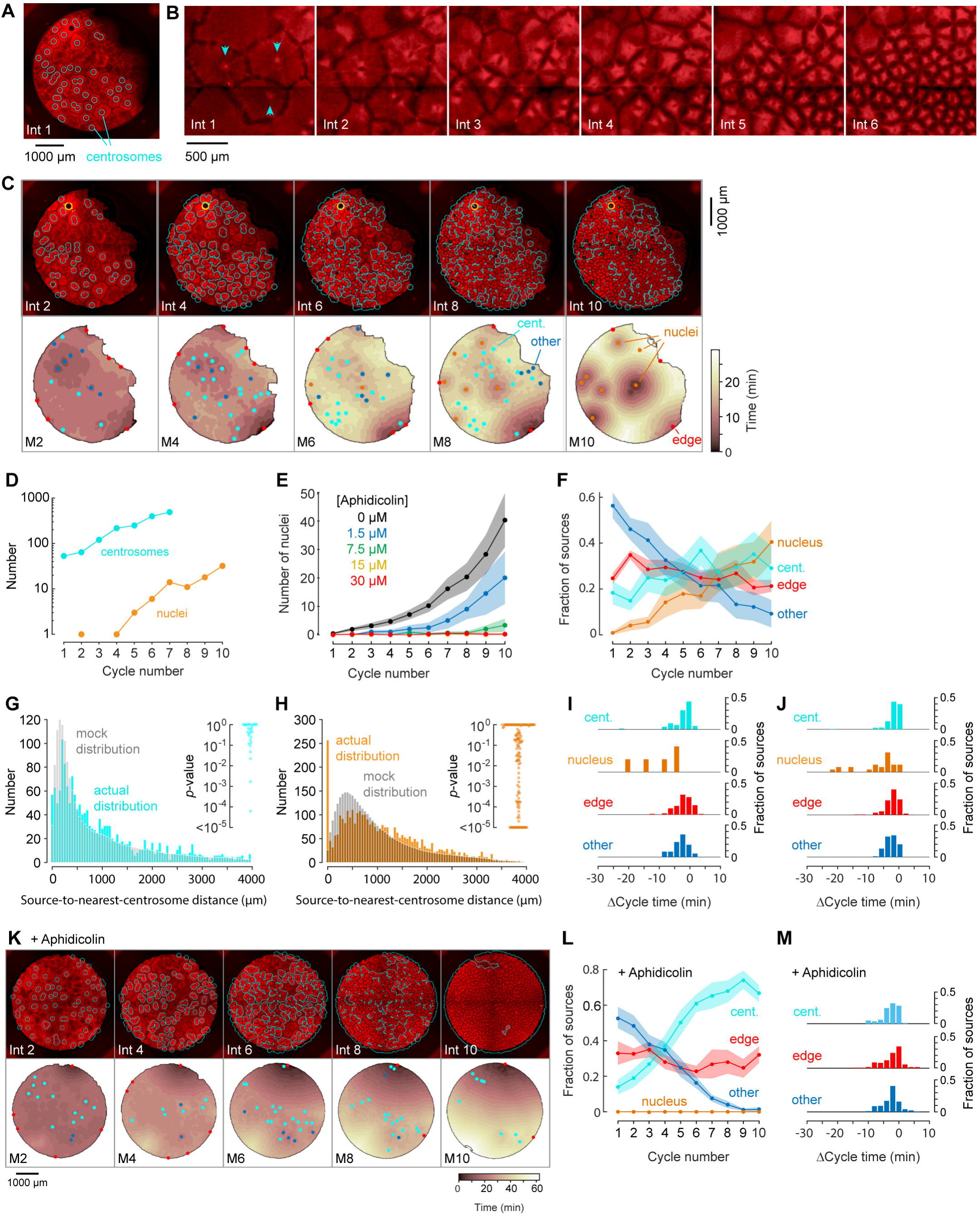
Centrosomes promote cytoplasmic organization and division of the cell-like compartments, but are not strong pacemaker sources. (A) Identification of centrosomes in an extract supplemented with purified HeLa cell centrosomes. (B) Montage showing that centrosomes duplicate and promote the division of cell-like compartments. SiR-Tubulin fluorescence is shown in red. (C) Montage showing SiR-tubulin fluorescence (red), NLS-mCherry fluorescence (green), and centrosome positions (cyan) (top), and heat maps of the mitotic sources (bottom), as a function of cycle number from a typical experiment. The coloring of the heat map denotes the time at which each pixel entered mitosis relative to the earliest pixels in that cycle. See also movie S6. (D) The number of centrosomes and adventiously-produced nuclei per cell cycle for the experiment shown in (C). (E) Suppression of nucleus formation by aphidicolin. (F) The fraction of the mitotic sources associated with nuclei/centrosomes (orange), edges (red), centrosomes (cyan) or others (blue) as a function of cycle number. The data points are mean fractions from 15 experiments where the extracts cycled at least 10 times, and the shaded regions show the ± SEM. (G, H) The distribution of distances from mitotic sources to the nearest centrosome (G, cyan), from 132 experiments, 18 cycles, and 1429 sources, and from mitotic sources to the nearest nucleus (H, orange), with same number of experiments and cycles as in G, but with 3595 sources. Only those cycles with fewer than 100 centrosomes or nuclei were included in G or H, respectively. The expected random distributions (mock distribution) of distances are shown in gray. The insets show probabilities (*p*-values) of obtaining the observed number of mitotic sources (or more than the observed number) that are close (< 100 µm) to centrosomes (G) or nuclei for the individual analyzed cycles. *p*-values were calculated by bootstrapping with 10^5^ randomized source positions for each individual cycle. (I) Acceleration of the cell cycle (measured as Δcycle times) relative to the slowest 15% of the cytoplasm. These Δcycle times were calculated for the first cell cycle after the appearance of centrosomes. Data are from 19 experiments and include 74 centrosome-, 5 nucleus-, 103 edge-associated sources and 167 other sources. (J) Acceleration of the cell cycle (measured as Δcycle times) relative to the slowest 15% of the cytoplasm, calculated for the first cell cycle after the appearance of nuclei (for those experiments where nuclei appeared). Data are from 18 experiments and include 258 centrosome-, 26 nucleus-, 515 edge-associated and 142 other sources. (K) Montage showing SiR-tubulin (red), NLS-mCherry (green), and centrosomes (cyan) for a centrosome-supplemented extract treated with 15 µM aphidicolin. (L) The fraction of the mitotic sources associated with centrosomes (cyan), edges (red) or others (blue) as a function of cycle number for aphidicolin-treated (15 to 60 µM), centrosome-supplemented extracts. From 11 experiments where the extracts cycled at least 10 times. (M) The Δcycle times for the three classes of mitotic sources. Data are from 14 experiments and include 82 centrosome-, 121 edge-associated sources and 169 other sources.

Nuclei sometimes eventually appeared in centrosome-supplemented extracts, even though no sperm chromatin had been added, and once they appeared, they replicated just as the centrosomes did (Fig. 4C, D). The appearance of nuclei was inhibited by the DNA polymerase inhibitor aphidicolin in a dose-dependent fashion (Fig. 4E), consistent with the hypothesis that they arose from the replication of small quantities of *Xenopus* egg DNA present in the egg extracts, or of HeLa cell DNA present in the purified centrosomes.

To determine whether centrosomes were acting as mitotic sources, we compared the distances between sources and the nearest centrosome to what would be expected by chance. The actual distribution of distances was similar to the mock distribution (Fig. 4G), and only 4/29 individual cycles had more sources within 100 µm of a centrosome than would be expected by chance (Fig. 4G, inset). Once nuclei appeared, some became trigger wave sources, and the number of sources close (< 100 µm) to a nucleus was substantially higher than what would be expected by chance (Fig. 4H; 61/132 of the individual cycles). As was the case with sperm-supplemented extracts, these nuclei accelerated the cell cycle more than centrosome-associated sources, edge-associated sources, or other sources did (Fig. 4I), and the proportion of sources associated with nuclei increased with cycle number (Fig. 4F).

Adding aphidicolin to centrosome-supplemented extracts did not alter the organization of the cytoplasm or the division of the cell-like compartments (Fig. 4J and movie S7), it but did suppress the appearance of nuclei and left the extract with mitotic sources that only modestly accelerated the cell cycle (Fig. 4K).

Thus, although HeLa cell centrosomes promoted microtubule organization and allowed the cell-like compartments to divide, they acted as relatively weak mitotic sources; they did not accelerate the cell cycle to a greater extent than edge-associated sources or sources not associated with particular structures.

### Phage DNA forms nuclei that act as robust trigger wave sources in the absence of centrosomes

To determine whether nuclei could substantially accelerate mitotic entry in the absence of centrosomes, we examined mitotic initiation in centrosome-less extracts that were supplemented with λ-bacteriophage DNA (Fig. 5A and movie S8). The phage DNA (nominally 5 µg/ml) formed nuclei, as previously reported (*32*), and the nuclei replicated, but generally the cell-like compartments did not divide. This indicates that centrosomes are required for compartment division and that the production of nuclei from the phage DNA did not result in the de novo production of centrosomes.

**Fig. 5.**
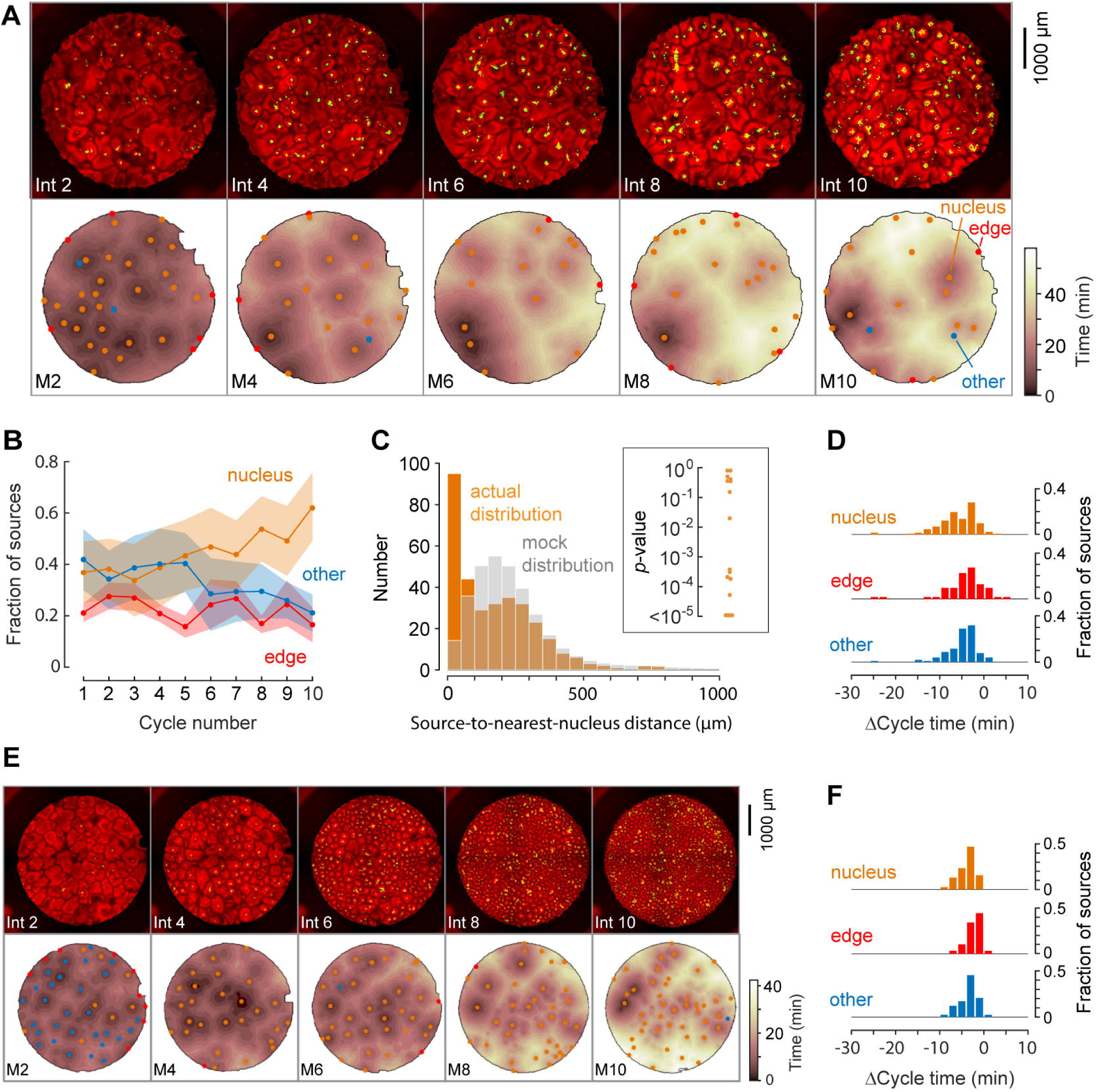
Phage DNA forms nuclei that act as trigger wave sources. (A) Montage from a typical experiment showing the microtubules (top) and the heat map representation of the microtubule sources (bottom) for cycles 2, 4, 6, 8, and 10. Nucleus-associated sources are shown in orange; edge-associated sources are shown in red; and other sources—i.e. those not associated with edges or nuclei—are shown in blue (other). The top panels show the extracts in interphase, with SiR-tubulin fluorescence in red and NLS-mCherry in green. The bottom panels are the heat maps of the mitotic sources. The coloring of the heat map denotes the time at which each pixel entered mitosis relative to the earliest pixels in that cycle. (B) Fraction of the mitotic sources associated with nuclei, edges, or no obvious structures as a function of cycle number. The data points are mean fractions from 4 experiments where the extracts cycled at least 10 times, and the shaded regions show the ± SEM. (C) The distance of the trigger wave sources to the nearest nucleus (orange), compared to a mock distribution (gray). Data are from 12 cell cycles from 4 experiments and include 235 sources. The inset shows probabilities (*p*-values) of obtaining the observed number of mitotic sources (or more than the observed number) that are close (< 100 µm) to nuclei for the 12 cell cycles individually, calculated by bootstrapping with 10^5^ randomized source positions for each individual cycle. (D) The duration of the cell cycle at nucleus-associated sources, edge sources, and other sources relative to the slowest 15% of the well. Data are from 8 experiments and include 130 nucleus-associated sources, 54 edge sources and 95 other sources). (E) Montage from an experiment with added phage DNA plus centrosomes. (F) The duration of the cell cycle at nucleus-associated sources, edge sources, and other sources relative to the slowest 15% of the well, for extracts with added phage DNA plus centrosomes. Data are from 3 experiments.

Many, but not all, of the nuclei became mitotic sources, and the initially strong sources became more dominant (deeper wells on the topographic map) with time (Fig. 5A). The fraction of sources contributed by nuclei increased with time (Fig. 5B), and the relative contributions of the nuclear, edge, and other sources was similar to that seen in sperm-supplemented extracts (Fig. 2E). Analysis of 12 cycles from 4 experiments showed that there were far more sources in proximity to nuclei than what would be expected by chance (Fig. 4C). Out of the 12 cycles, 9 had more sources in close proximity (< 100 µm) to nuclei than would be expected by chance (*p* < 0.05).

We assessed the cell cycle acceleration produced by the phage DNA by comparing the cycle times at the nuclear sources to the slowest 15% of pixels in the first cycle where nuclei were apparent. The distribution of Δcycle times for the three classes of sources is shown in Fig. 5D. The average acceleration at nuclear sources was 6.9 ± 4.5 min (Fig. 5D), about half what was seen with nuclear sources in sperm-supplemented extracts (Fig. 2I), but higher than what was typically seen for edge sources, centrosome-associated sources, or other sources (Figs. 2-4). The acceleration seen in the phage DNA experiments may be an underestimate; the nuclear sources initially present in the phage-supplemented extracts were so numerous that little if any of the extract appeared to not be entrained by a trigger wave. Alternatively, it is possible that nuclei formed around phage DNA have somewhat slower intrinsic rhythms than nuclei formed around sperm chromatin, or that nuclei lacking centrosomes have slower rhythms than nuclei that possess them.

To test this third possibility, we incubated extracts with purified HeLa cell centrosomes plus phage DNA and assessed mitotic dynamics. As shown in Fig. 5E, the centrosomes facilitated the organization of the extract and allowed the cell-like compartments to divide, as expected. However, there was no obvious increase in the acceleration of the cell cycle (Fig. 5F) as measured by the Δcycle times in three experiments. Thus if there an additional effect of having centrosomes present as a staging platform for mitosis, it is too subtle for this type of analysis to detect.

## Discussion

Here we used *Xenopus* egg extracts and reconstitution to determine what structure acts as pacemaker for the mitotic oscillator. We found that nuclei that were formed from *Xenopus* sperm chromatin (Figs. 1 and 2) or phage DNA (Fig. 5) acted as robust sources of mitotic trigger waves. This could be attributed to an acceleration of the cell cycle by about 10 min. In the absence of nuclei, added centrosomes facilitated the organization of cell-like compartments and allowed the cell-like compartments to divide, but had little ability to act as trigger wave sources (Fig. 4).

These findings support the hypothesis that the nucleus normally serves as the cell cycle pacemaker. It seems plausible that the nucleus’s ability to concentrate Cdc25C and cyclin B-Cdk1 (*21, 24, 26*) could be the mechanistic basis for the pacemaking, although a direct experimental test of this hypothesis has not yet been carried out.

These studies underscore that diverse biological processes, for example heart beats and cell cycles, may occur on different time scales (∼1 sec vs. ∼40 min) and involve different classes of proteins (ion channels vs. kinases, phosphatases, and ubiquitin ligases), but share an underlying unity in terms of their systems-level logic. In both cases the rhythms are generated by a relaxation oscillator circuit with a bistable rhythm, and in both cases this allows the rhythm to spread in an orderly fashion from a spatially-distinct pacemaker locus.

While this manuscript was in preparation, Nolet and colleagues published a paper that also argues that nuclei function as cell cycle pacemakers in *Xenopus* extracts, based on a combination of theory and experiments in a one-dimensional extract system (*33*). Our work agrees, complements, and extends their findings by directly assessing the contribution of the centrosome to the pacemaker function.

## Acknowledgments

We thank Yuping Chen and Shixuan Liu for comments on the manuscript, and the Ferrell and Stearns labs for helpful discussions. This work was supported by grants from the National Institutes of Health (T32 GM007276-43, R01 GM110564, R35 GM131792, and R35 GM120286).

## Author contributions

Conceptualization: O.A. and J.E.F.; Formal analysis: O.A.; Funding acquisition: J.E.F. and T.S.; Investigation: O.A. and G.K.B.; Supervision: J.E.F. and T.S.; Visualization: O.A. and J.E.F.; Writing: O.A. and J.E.F.; Editing: O.A., G.K.B., J.E.F and T.S.

## Data and materials availability

All data, code, and materials used in the analysis will be provided to any researcher for purposes of reproducing or extending the analysis.

