## Supplementary Materials for "The nucleus serves as the pacemaker for the cell cycle"

### Methods

### Figures S1-S2

### Figure S1. Montage of dividing cell-like compartments in an extract with no added sperm or centrosomes. Related to Fig. 3.

**Figure S2.** Montage of an extract with added boiled centrosomes (A) or normal centrosomes (B). Note there appears to be one functional centrosome in interphase 1 in (A), which subsequently replicates and divides, versus numerous functional centrosomes in (B).

**Movies S1-S8**

**Movie S1.** Trigger waves of tubulin depolymerization and polymerization in *Xenopus* extract supplemented with sperm. Related to Fig. 1B-F.

**Movie S2.** Three classes of trigger wave sources: others (left), nucleus-associated (center), and edge-associated (right). Related to Fig. 1G-I.

**Movie S3.** Close up of a dividing nuclear trigger wave source. Related to Fig. 1.

**Movie S4.** Trigger waves of tubulin depolymerization and polymerization in a second *Xenopus* extract supplemented with sperm. Related to Fig. 2.

**Movie S5.** Trigger waves of tubulin depolymerization and polymerization in a *Xenopus* extract supplemented with no sperm, centrosomes, or added nuclei. Related to Fig. 3.

**Movie S6.** Trigger waves of tubulin depolymerization and polymerization in a *Xenopus* extract supplemented with purified HeLa cell centrosomes. Related to Fig. 4.

**Movie S7.** Trigger waves of tubulin depolymerization and polymerization in a *Xenopus* extract supplemented with purified HeLa cell centrosomes and aphidicolin (15 µM). Related to Fig. 4.

**Movie S8.** Trigger waves of tubulin depolymerization and polymerization in a *Xenopus* extract supplemented with λ-bacteriophage DNA (5 µg/mL). Related to Fig. 5.

**Methods**

#### Extract preparation

Frog handling and egg extract preparation is as described in (*36*), except that the extract was typically clarified for 3 times for 8 min at 16,000 g to improve optical transparency. To prevent the over-dilution of the extract, staining, addition of components or drugs, was typically done at a ratio of 1:100. SiR-Tubulin (Spirochrome, dissolved in DMSO) was used for staining at 0.3 μM (0.3% final DMSO concentration). Aphidicolin (Calbiochem; dissolved in DMSO) was applied as indicated in text (0.16% final concentration of DMSO for every 10 μM of aphidicolin). GST-NLS-mCherry was purified and used as described in the Methods section of reference (*16*). λ-DNA (New England Biolabs) was used at 0.5 μg per 100 μl of extract. Centrosomes were purified following (*37*), and were typically applied at 30-1000 centrosomes per 100 μl of extract. Demembranated sperm was prepared as described (*30*), and was typically used at 30-1000 sperm counts per 100 μl of extract.

#### Imaging

For imaging, 3 μl of extract was spread on the bottom of a well of a Corning 96-well polystyrene plate (#3368), and covered with 200 μl of heavy mineral oil (Sigma). The extract was imaged in time-lapse using a Leica epifluorescence microscope and a 5x objective. Acquisition frequency varied between 30 s to 2 min per frame.

#### Image analysis

Images were automatically stitched, segmented, and analyzed using a custom-made pipeline scripted in Matlab. Code is available upon request. Mitotic sources and nuclei were located automatically and were checked manually. Centrosome positions were determined manually. Because the initiation time for the interphase of the first cycle was unknown (i.e., the segment of time before the first mitosis), all of the analysis preformed starting from the first mitosis.

#### Bootstrapping calculation

Bootstrapping was performed by randomizing the locations of mitotic initiation for a cycle, determining how many of these randomized sources were within 100 µm of a nucleus or a centrosome, and then repeating the procedure 100,000 times. The *p*-value for that cycle was calculated as the number of iterations in which there were as many or more sources in proximity by random, divided by 100000.

**Figure S1**

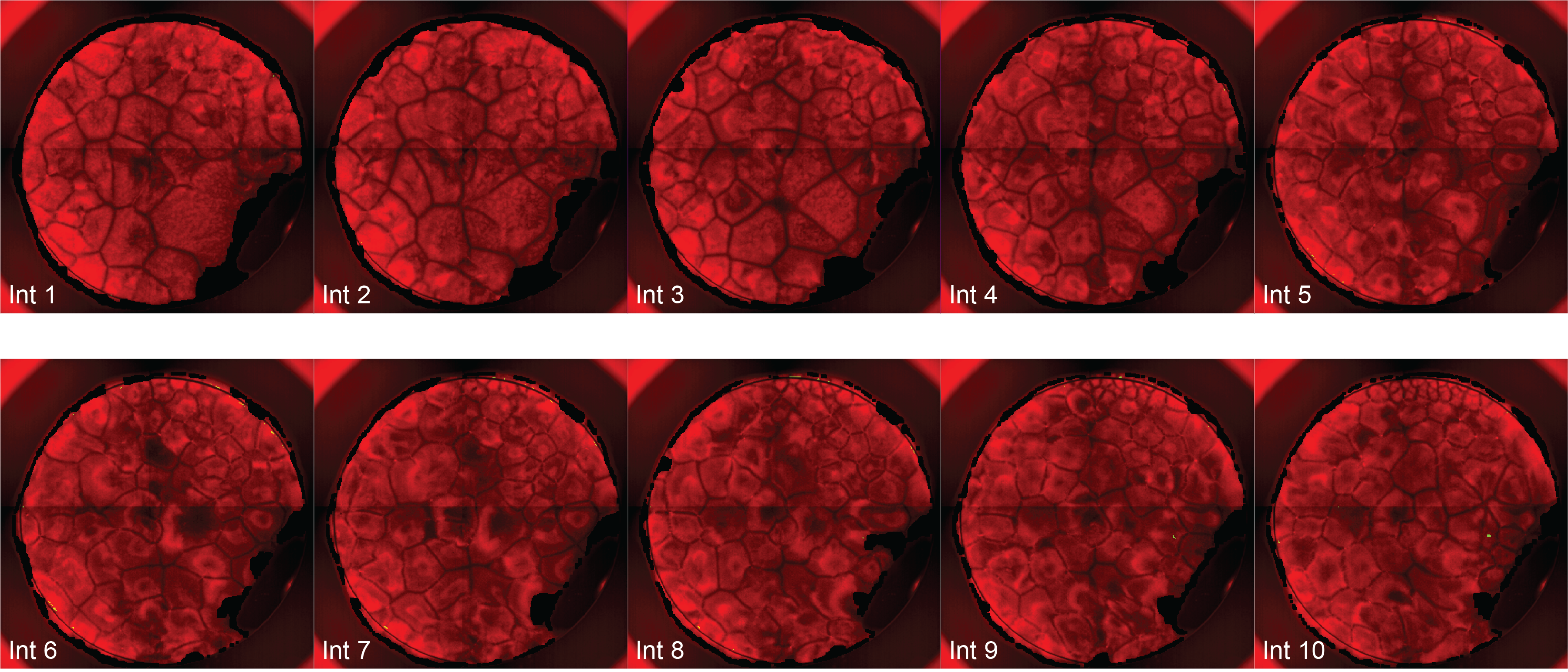

**Fig, S1. Emergence of dividing cell-like compartments near the edge of an extract not supplemented with sperm, centrosomes, or phage DNA.** From cycle 6 on, the cell-like compartments at ~12:00 divide with each cycle.

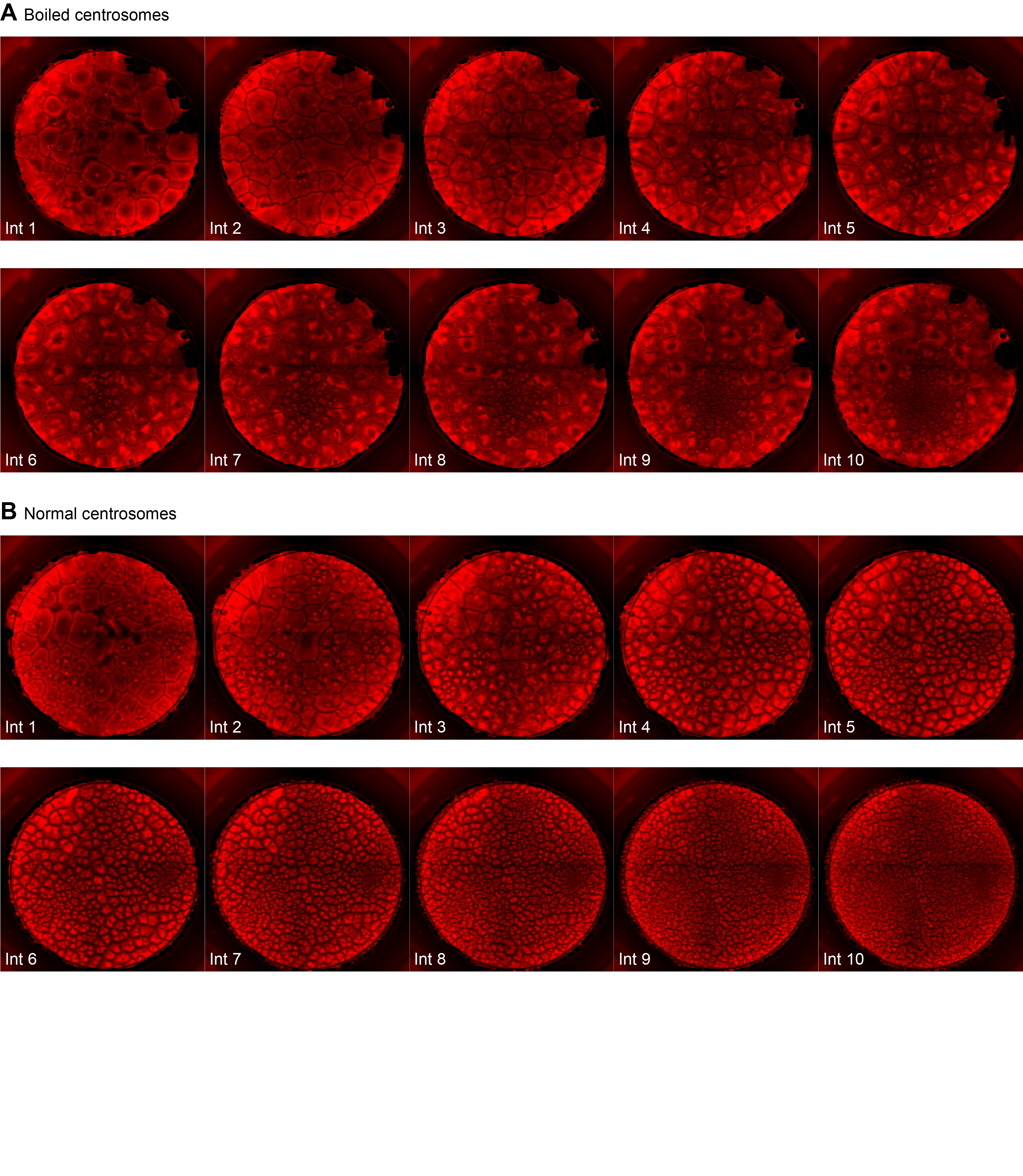

**Figure S2.** Montage of an extract with added boiled centrosomes (A) or normal centrosomes (B). Note there appears to be one functional centrosome in interphase 1 in (A), which subsequently replicates and divides, versus numerous functional centrosomes in (B).
